# Single-shot super-resolution total internal reflection fluorescence microscopy

**DOI:** 10.1101/182121

**Authors:** Min Guo, Panagiotis Chandris, John Paul Giannini, Adam J. Trexler, Robert Fischer, Jiji Chen, Harshad D. Vishwasrao, Ivan Rey-Suarez, Yicong Wu, Clare M. Waterman, George H. Patterson, Arpita Upadhyaya, Justin Taraska, Hari Shroff

## Abstract

We demonstrate a simple method for combining instant structured illumination microscopy (SIM) with total internal reflection fluorescence microscopy (TIRF), doubling the spatial resolution of TIRF (down to 115 +/-13 nm) and enabling imaging frame rates up to 100 Hz over hundreds of time points. We apply instant TIRF-SIM to multiple live samples, achieving rapid, high contrast super-resolution imaging in close proximity to the coverslip surface.

---

Total internal reflection fluorescence (TIRF) microscopy^1^ provides unparalleled optical sectioning, exploiting an evanescent field induced at the boundary between high and low refractive index media to selectively excite only fluorophores within one wavelength of the coverslip surface. The superb background rejection, low phototoxicity, high speed, and sensitivity of TIRF microscopy has been used to study diverse biological phenomena at the plasma membrane, including endocytosis, exocytosis, and focal adhesion dynamics. TIRF microscopy has also been combined with super-resolution methods, particularly structured illumination microscopy (SIM^2-5^) to enable subdiffractive imaging in living cells^3,6^. Unfortunately, all previous methods sacrifice temporal resolution to improve spatial resolution, limiting the effectiveness of TIRF in studying dynamic phenomena.

We and others have developed SIM implementations that improve spatial resolution without compromising speed^7-9^. These microscopes sharpen the image ‘instantly’ (i.e. during image formation) by optically combining information from excitation- and emission-point-spread functions (PSFs), obviating the need to acquire and process extra diffraction-limited images that slows classic SIM. our previous instant SIM design^7,10^ modified a swept field confocal geometry, scanning an array of sharp excitation foci to elicit fluorescence, de-scanning the fluorescence, rejecting out-of-focus fluorescence with a pinhole array, and locally contracting each focus before rescanning to produce a super-resolution image. Motivated by our success in using instant SIM for high speed super-resolution imaging, we sought to adapt the same underlying concept for TIRF.

TIRF is enabled when highly inclined light with incidence angle θ≥= θ_C_ = arcsin (n_2_/n_1_) impinges upon the boundary between media with indices n_1_ and n_2_, with n_1_ > n_2_. We reasoned that placing an annular mask at a Fourier image plane (optically conjugate to the back focal plane of the objective) would block all subcritical rays, thereby producing TIRF without otherwise perturbing the speed and functionality of our original instant SIM. Annular illumination has been used to generate a single TIRF spot in diffraction-limited^11^ and stimulated emission depletion microscopy^12^, yet for parallelized instant SIM an array of spots is needed.

We created such a pattern by carefully positioning an annulus one focal length away from the foci produced by our excitation microlens array, thus simultaneously filtering out low angle rays in each excitation focus. The resulting beams were relayed to the sample by instant SIM optical components, including a two-sided galvanometric mirror conjugate to the back focal plane of the objective (a 1.7 numerical aperture (NA) lens used for the large range of accessible θ≥θC, facilitating TIRF). Emission optics were nearly identical to the original instant SIM setup (**Methods**), and included pinhole- and emission microlens arrays with appropriate relay optics (**Supplementary Fig. 1**).

**Figure 1.**
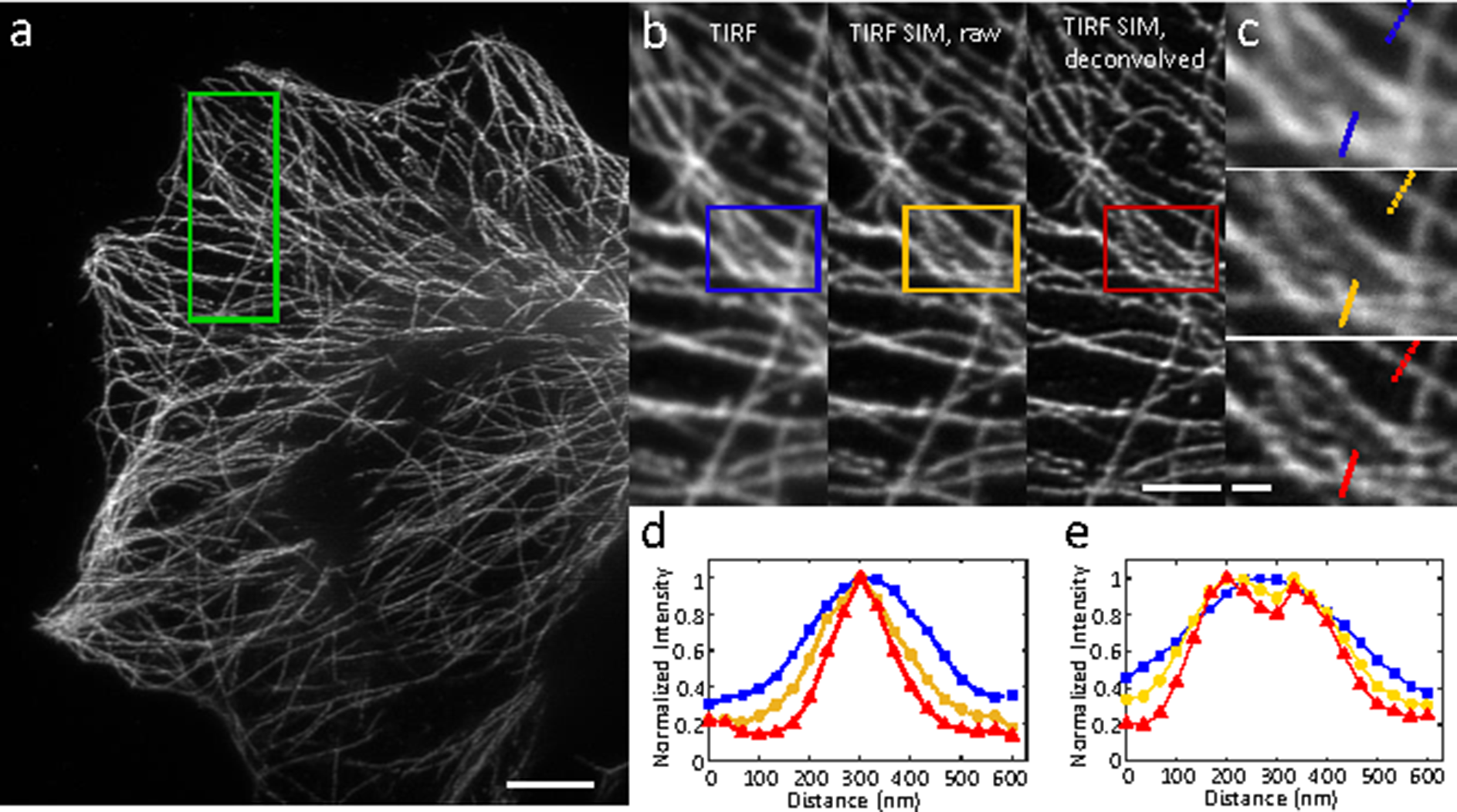
Resolution enhancement via instant TIRF-SIM. **a)** Deconvolved instant TIRF-SIM image of immunolabeled microtubules in a fixed U2oS cell. **b)** Higher magnification views of the green rectangular region in **a)** showing diffraction limited TIRF (obtained using only the excitation microlenses, left), instant SIM (raw data after employing pinholes and emission microlenses, middle), and deconvolved instant SIM (right). **c)** Higher magnification views of TIRF (top), raw instant SIM (middle) and deconvolved instant SIM images, corresponding to blue, yellow, and red rectangular regions in **b).** Comparative line profiles **(d**, dashed lines in **c**; **e**, solid lines in **c**) are also shown. Scale bars: 5 μm in **a**, 2μm in **b**, 0.5 μm in **c**.

Since annular excitation produces a focused spot with pronounced sidelobes (due to the Bessel-like character of the excitation), we were concerned that interference between neighboring foci and transfer of energy from the central intensity maxima to the sidelobes would significantly diminish illumination contrast in the focal plane (**Supplementary Note 1**). Indeed, during imaging of fluorescent dye in TIRF mode, we did observe substantial background fluorescence between excitation foci (yet still less than observed when imaging conventionally, due to the dramatic reduction of out-of-focus fluorescence in TIRF). However, individual foci were sharply defined and the extraneous background could be readily removed with the pinhole array intrinsic to our setup (**Supplementary Fig. 2**). We confirmed that TIRF was maintained during the imaging process by measuring the depth of the evanescent field with index-matched silica beads (**Supplementary Fig. 3**), finding this value to be 123 nm+/-6 nm (95% confidence interval). Qualitative comparisons on fixed microtubule samples also demonstrated the improved sectioning in TIRF, relative to conventional instant SIM **(Supplementary Fig. 4)**.

**Fig. 3.**
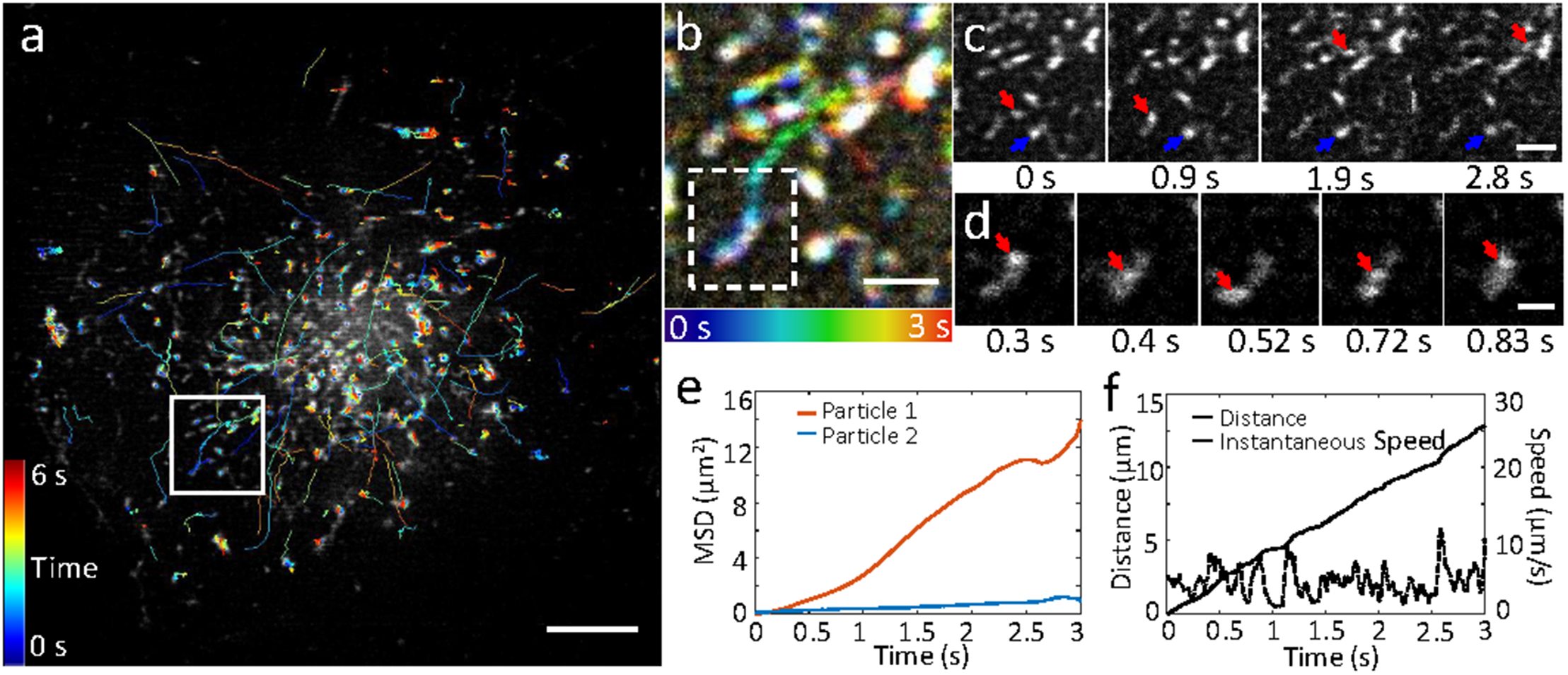
Rapid dynamics of Rab11 are resolved at 100 Hz with instant TIRF-SIM. EGFP-Rab11 was transfected into U2oS cells and imaged at 37°C at 100 Hz. **a)** First frame from image series, with overlaid tracks (lines colored to indicate time, color bar indicated at left). **b)** Higher magnification view of white rectangular region in **a)**, over the first 3 seconds of acquisition. Time evolution indicated in color according to bar at bottom. **c)** Selected raw images corresponding to imaging region **b)**, emphasizing motile (red arrow) and more stationary (blue arrow) particle. **d)** Magnified view of more motile particle indicated with red arrow in **c)**, highlighting bidirectional motion. Mean square displacements of both particles (**e**) and distance and instantaneous speed (**f**) of the motile particle are also quantified. Scale bars: 5 μm in **a**; 1 μm in **b**, **c**; 0.5 μm in **d**. See also **Supplementary Figs. 10** and **Supplementary Videos 10, 11**.

**Figure 2.**
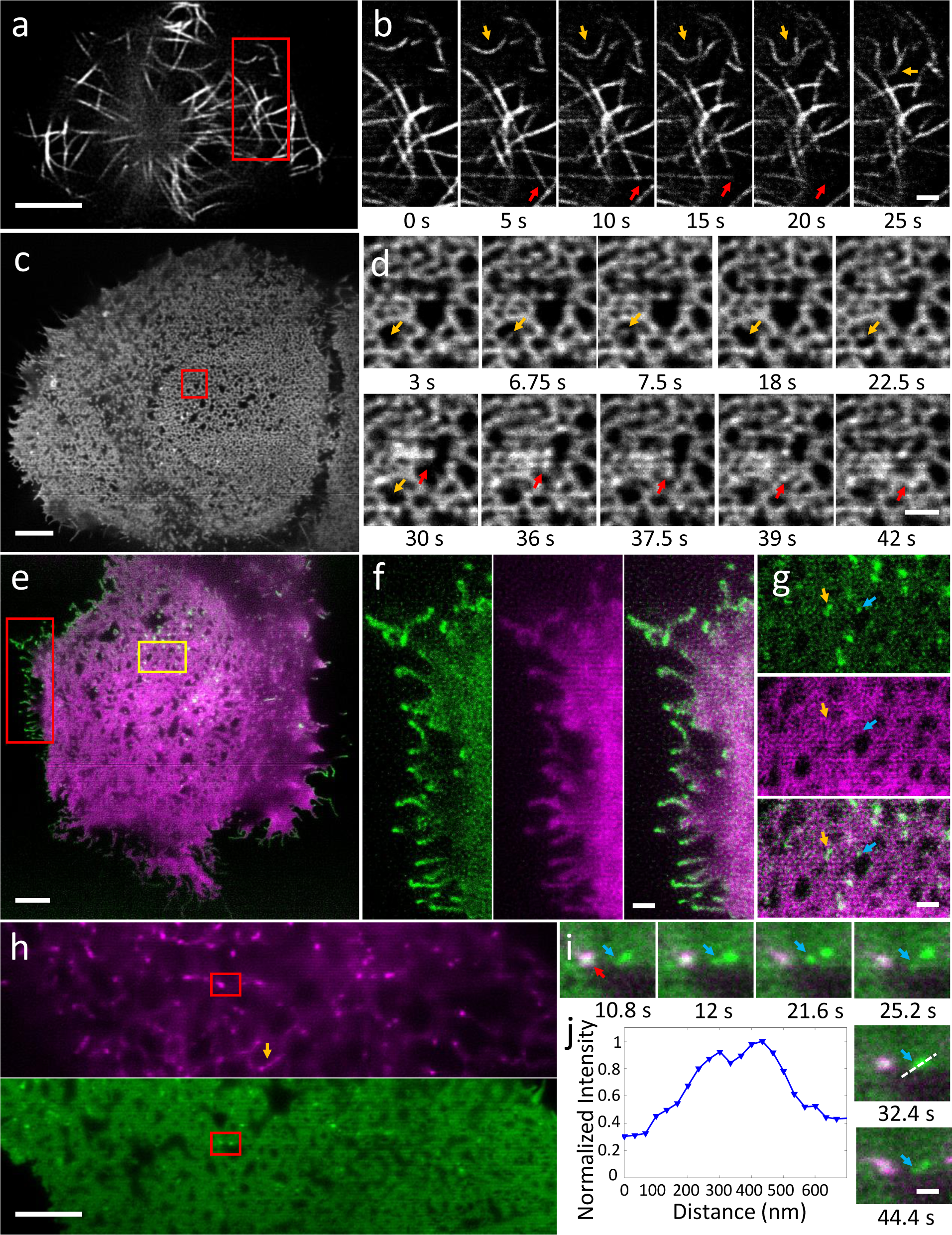
Instant TIRF-SIM enables high speed super-resolution imaging at the plasma membrane over hundreds of time points. **a**) Image of EMTB-3xEGFP expressed in Jurkat T cells, taken from 500 frame series (images recorded every 50 ms). Higher magnification series **b)** of red rectangular region in **a** highlights microtubule buckling (orange arrows) and movement (red arrows mark two microtubule bundles that move left and up during image series). See also **Supplementary Video 1**. **c)** Image of EGFP-HRAS expressed in U2oS cell, taken from series spanning 60 time points, images recorded every 0.75 s.**d)** Higher magnification view of red rectangular region in **c)** emphasizing dynamics, including transient filling in (orange arrows) and reorganization (red arrows) of microdomains. See also **Supplementary Videos 2, 3**. **e)** Two-color image showing EGFP-VSVG (green) and Halotag-Ras (labeled with Janelia Fluor 546, magenta), derived from series spanning 100 time points, dual-color images recorded every 2.3 s. **f)** Higher magnification view of red rectangular region in **e)**, showing EGFP-VSVG (left), Halotag-Ras (middle) and merged (right) distributions, highlighting concentrated VSVG at cell periphery. **g)** Higher magnification view of orange rectangular region in **e**), showing EGFP-VSVG (top), Halotag-Ras (middle) and merge (bottom). Arrows mark VSVG puncta located near Ras microdomains. See also **Supplementary Video 4**. **h)** Two-color image showing pDsRed2 ER (magenta, top) and EGFP-HRas (green, bottom), derived from image series spanning 100 time points, images recorded every 1.2 s. orange arrow highlights ER tubule. **i)** Higher magnification series of red rectangular region in **h**, highlighting dynamics of Ras puncta (blue arrow) in vicinity of ER contact site (red arrow). **j)** profile of dashed line in **i** indicating peak-to-peak separation of 134 nm in HRas channel. See also **Supplementary Video 5, 6**. Scale bars: 5 μm in **a, c, e, h**; 1 μm in **b, d, f, g**; 500 nm in **i**.

We estimated system resolution on 100 nm fluorescent beads (**Supplementary Fig. 5**). With scanned TIRF excitation only, beads were resolved to 249 +/-11 nm (N = 20 beads, mean +/-standard deviation). Descanning, pinholing, locally contracting, and rescanning reduced the apparent bead diameter to 194 +/-20 nm, and resolution could be further improved after deconvolution (10 iterations, Richardson-Lucy deconvolution) to 115 +/-13 nm. These results were similar to those obtained using conventional illumination with the same objective lens, i.e. with conventional instant SIM (**Supplementary Table 1**) implying that our spatial resolution did not degrade with TIRF. Images of fixed cells further confirmed this progressive resolution improvement **(Fig. 1a-c)**, as individual microtubules had an apparent width of ˜128 nm (**Fig. 1d**), and we were able to distinguish microtubules spaced 134 nm apart (**Fig. 1e**). Qualitative tests in living cells confirmed our resolving power (**Supplementary Fig. 6**), as we were able to observe individual GFP-labeled myosin IIA heads^13^ and void areas within GFP-FCHo2 puncta^14^, subdiffractive structural features that have previously been resolved with TIRF-SIM.

We next used instant TIRF-SIM to examine the dynamics of protein distributions in living cells (**Fig. 2**). First, we recorded microtubule dynamics over 500 time-points by imaging the fluorescence microtubule binding probe, EMTB-3xEGFP^15,16^, in Jurkat T cells after they settled on anti-CD3 coated coverslips (**Fig. 2a**, **Supplementary Video 1**). our imaging rate of 20 Hz was sufficient to easily follow buckling, shortening, and sliding of microtubule bundles at the base of the cell within the evanescent TIRF field (**Fig. 2b**). As a second example, we recorded the dynamics of the small GTPase H-Ras, which is lipidated and then targeted to the plasma membrane^17^. Images were acquired every 0.75 s over 100 timepoints in U2oS cells (**Fig. 2c**, **Supplementary Video 2**). Intriguingly, GFP-HRas localized in highly dynamic microdomains at the plasma membrane (**Fig. 2c, d, Supplementary Video 3**). The high spatiotemporal resolution of our technique revealed rich dynamics of this reticulated pattern, as we observed reorganization of domains with a temporal resolution of about 1.5 s, including transient ‘filling in’of the void areas between microdomains (**Fig. 2d**), and coordinated, ‘wave-like’motion between microdomains (**Supplementary Video 3**). To our knowledge, neither the distribution nor the dynamics of Ras has been reported at this length scale in living cells, perhaps due to the lack of spatial resolution or optical sectioning (e.g., we found that in diffraction-limited TIRF imaging or conventional instant SIM at lower NA, microdomains were poorly resolved, **Supplementary Fig. 7**).

We also imaged Halotag-HRas (labeled with Janelia Fluor 549^18^) in combination with GFP-tagged vesicular stomatitis virus G protein (VSVG, **Fig. 2e, Supplementary Video 4**), highlighting the ability of instant TIRF-SIM for live, dual-color imaging at the plasma membrane. VSVG traffics from the endoplasmic reticulum (ER)-Golgi system to the plasma membrane in a vesicular manner marking the secretory pathway, but also shuttles back to the Golgi system using the endosomal compartment as a carrier^19^. Despite similar targeting to the plasma membrane^20^, GFP-VSVG and Halotag-HRas displayed distinct localization within living cells (**Fig. 2g**). VSVG showed some localization around Ras microdomains within the cell interior (**Fig. 2g**) but we also observed preferential enrichment of VSVG at the cell boundary, particularly at cell filopodia and filamentous structures. In a second dual-color example, we imaged GFP-HRAS with pDsRed2-ER **(Fig. 2h**), which carries the KDEL luminal ER retention signal sequence and marks the endoplasmic reticulum. Given the optical sectioning of our technique,the ER mostly appeared as a set of bright punctate spots and occasional tubules in close proximity to the plasma membrane, while the rest of the ER appeared as a network structure presumably further from the coverslip. Although punctate ER structures occasionally colocalized with Ras, the protein distributions were mostly distinct and exhibited different dynamics (**Supplementary Video 5**), consistent with their differential localization and function within the cell. The spatial resolution of our technique proved key in resolving apparent fission and fusion of Ras microclusters adjacent to more stable ER contacts (**Fig. 2i, Supplementary Video 6**), a phenomenon otherwise obscured by diffraction (**Fig. 2j**).

We also visualized intracellular calcium flux (**Supplementary Video 7**), actin (**Supplementary Video 8, 9**), and myosin IIB dynamics (**Supplementary Video 10**) in live cells. These examples all underscore the ability of instant TIRF-SIM to enable super-resolution imaging well matched to the dynamics of interest, either matching or surpassing the image acquisition rate offered by standard TIRF-SIM systems (**Supplementary Table 2**).

A key advantage in instant SIM is the ability to image at much faster frame rates, because the super-resolution image is formed during a single camera exposure. To illustrate this capability, we imaged GFP tagged Rab11, a recycling-endosome specific GTPase that drives constant turnover of endosomes from the plasma membrane to the cytosol and modulates extracellular release of vesicles^21^, in U2oS cells at 37°C at 100 Hz (**Supplementary Video 11**). This imaging rate was sufficient to visualize and track the rapid motion of 980 Rab11-decorated particles (**Fig. 3a**). An analysis of track motion revealed that the majority of particles underwent < 1 **µ**m displacement over our 6 s imaging period, yet we also observed tens of particles that showed greater displacements (**Supplementary Fig. 8**). Particles that traveled farther also traveled faster, with mean speed greater than 1 μm/s (**Supplementary Fig. 8**) and with instantaneous speed in some cases exceeding 10 **μ**m/s (**Fig. 3d, f**). A closer analysis at the single particle level (**Fig. 3b, c**) also revealed qualitative differences in particle motion, with some particles undergoing diffusive motion, as revealed by a linear mean square displacement (MSD) vs. time and others showing supralinear MSD vs. time (**Fig. 3e**) with bouts of directed motion (**Fig. 3d**, **Supplementary Video 12, 13**). We note that imaging at slower frame rates (as with imaging with any previous implementation of TIRF-SIM) distorts track lengths because long tracks are broken into shorter tracks, short tracks are discarded, and multiple independent short tracks may be classified falsely as longer tracks (**Supplementary Fig. 9**).

Our instrument provides fundamentally faster operation than classic TIRF-SIM systems, as only one image needs to be acquired, instead of the standard nine^3^. Additional advantages of our implementation over alternative approaches include less read noise (since fewer images are acquired) and less computational processing (our method requires only simple deconvolution of the raw images, instead of extensive image processing in Fourier space). Although the spatial resolution we report (˜115 nm) is ˜40% worse than claimed in state of the art linear TIRF-SIM^6^ (84 nm), our existing implementation of instant TIRF-SIM is ˜50 fold faster, as we demonstrate by imaging at frame rates up to 100 Hz (instead of ˜ 2 Hz).

Room for technical improvement remains. The excitation efficiency of our setup is low, as ˜60% of the illumination is blocked by the annular mask. Using a spatial light modulator (SLM) to generate the pattern might direct the illumination through the annular mask much more effectively, allowing lower power lasers to be used. Control over the phase of the illumination might also reduce the sidelobes in each of the foci, improving contrast in the focal plane and perhaps even removing the need for pinholes (although we suspect that pinholes are still useful in reducing scattered light that continues to plague objective-based TIRF^22^). Using optics that allow rapid adjustment of the annulus dimensions (such as an SLM or a digital micromirror device) might enable easy adjustment of the evanescent field depth, thereby providing additional axial information within the TIRF zone^23^. Finally, we did not exploit the narrower central maximum in each excitation focus for (marginally) higher spatial resolution, due to the coupling between inter-focus distance and focus size (**Supplementary Note 1**). Combining TIRF with single-point rescanning SIM^9,24^ would address this issue due to the more flexible design, albeit at the cost of temporal resolution.

## Acknowledgements

We thank Steve Lee and Abhishek Kumar for useful discussion on the instant TIRF-SIM, Henry Eden and Patrick La Riviere for providing feedback on the manuscript, Chris Combs for loaning us the Leica alignment reticle, Ethan Tyler for help with the illustrations, William Bement for the gift of EMTB-3xEGFP, Dyche Mullins for the gift of pEGFP-C1 F-tractin-EGFP, Dominic Esposito for the gift of HaloTag chimera of HRas, and Luke Lavis for the gift of Janelia Fluor549. Support for this work was provided by the Intramural Research Programs of the National Institute of Biomedical Imaging and Bioengineering; the National Heart, Lung, and Blood Institute; A.U. and I.R. were supported by NSF grant 1607645. JT and HS acknowledge funding from the NIH Director ‚s Challenge Innovation Award Program.

## Author contributions

Conceived project: J.T. and H.S. Designed optical layout: M.G., J.G., and H.S. Built optical system: M.G. and J.G. Acquired data: M.G., P. C. and J. C. Performed simulations: M.G. and Y.W. Prepared biological samples: P.C., A.J.T., R.F., J.C., H.V., and I.R-S. Provided advice on biological samples: P.C., R.F., C.W., A.U., and J.T. Performed tracking analysis: M.G. and J.C. with advice from I.R.-S. Provided expert advice on TIRF: G.H.P., J.T. Wrote paper: M.G. and H.S. with input from all authors. Supervised research: H.S.

## Methods

### Instant TIRF-SIM

The instant TIRF-SIM is built directly upon our previously reported instant SIM system^7^, but with two important modifications in the excitation path. First, we used a 1.7 NA objective (olympus, APoN100XHoTIRF) for excitation and detection. When imaging into aqueous samples with refractive index 1.33, 1-(1.33/1.7) = 0.22 of the objective back focal plane diameter (*d*_BFP_) is available for TIRF, implying that sub-critical illumination rays within a diameter 0.78 * *d*_BFP_ = 0.78 * 2 * NA_oBJ_ * *f*_oBJ_ = 0.78 * 2 * 1.7 * 1.8 mm = 4.77 mm must be blocked. Second, we inserted a relay system into the excitation arm of the instant SIM to block these rays. Excitation from 488 nm and 561 nm lasers was combined and beam expanded as before, and directed to a microlens array (Amus, f = 6 mm, 222 mm spacing between microlenses, 1 mm thick, 25 mm diameter, antireflection coated over 400–650 nm, APo-Q-P222-F6(633)+CHR) to produce an array of excitation foci. We used a matched pair of scan lenses (Scan lens 1 and 2, f=190 mm, Special optics, 55-S190-60-VIS) placed in 4*f* configuration to relay these excitation foci to the rest of the optical system, inserting an opaque circular mask (Photosciences, 2.68 mm diameter chrome circle with optical density 5 on 4″ x 4″ x 0.090″ quartz wafer) at the focal point between scan lenses (and the Fourier plane of the excitation foci produced by the microlens array) to filter subcritical rays. Given the 350 mm/ 190 mm = 1.84x magnification between the mask and the back focal plane of the objective, we designed the mask to block the central 2.68 mm * 1.84 = 4.93 mm diameter of the illumination. An iris placed just after the mask ensured that the outer diameter of the beam was ˜3.33 mm, a diameter that magnified to 3.33 * 1.84 = 6.13 mm, or ˜*d***_BFP_**, thereby reducing stray light that would otherwise fall outside the objective back focal plane. Alignment of the opaque mask and microlens array, which is critical, was greatly aided by placing the former on a 3-axis translation stage (Thorlabs, LT3, used for correct positioning of the mask image at the back focal plane) and the latter on a uniaxial translation stage (Thorlabs, LNR50M, used to position excitation foci precisely at the focal plane of the objective lens). We also used an alignment reticle (Leica) that screwed into our objective turret to further check that the annular illumination pattern was properly positioned (concentric with the optical axis of the objective) and focused at the back focal plane of our objective.

In the emission path, optics were identical to our previous design, except that we used a pinhole array with larger pinholes (Photosciences, Chrome on 0.090″ thick quartz, 222 μm pinhole spacing, 50 μm pinhole diameter) and an emission-side microlens array with longer focal length (f = 1.86 mm, Amus, APo-Q-P222-F1.86(633)). The total magnification between sample and our scientific grade complementary metal-oxide semiconductor camera (PCo-TECH, pco.edge 4.2) detector was 350 mm /1.8 mm = 194.4, resulting in an image pixel size of 33.4 nm. These elements are shown in **Suplementary Fig. 1**.

The excitation laser power was measured immediately prior to the objective. Depending on the sample, the average power ranged from 0.2 −2 mW, implying an intensity range from ˜7 – 70 W/cm^2^ (given our 58 μm x 52 μm field of view).

Samples were deposited on 20 mm diameter high index coverslips (olympus, 9-U992) designed for use with the 1.7 NA lens. Coverslips were mounted in a magnetic chamber (Live Cell Instrument, CM-B20-1) that attached to the microscope stage. For temperature maintenance at 37 °C, the magnetic chamber was mounted within an incubation chamber (okolab, H301-MINI).

### Estimating the evanescent field depth

We used two methods to estimate evanescent field depth. First, we used an analytical method^25^. For excitation of wavelength λ impinging at angle θ_1_ upon an interface with indices n_1_ and n_2_, n_1_ µ n_2_, the intensity I of an evanescent field decays along the optical axis with decay constant d according to I(z) = I_o_exp(-z/d), with 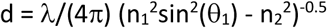. The term 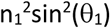 is equivalent to the square of an “ effective ” NA, in our case ≤ 1.7. If considering the smallest angles in our annular excitation (corresponding to the inner radius used in the mask, and producing evanescent waves with the longest decay length), this effective NA is 4.93/6.12 * NA_oBJ_ = 1.37. Assuming n_2_ = 1.33 and λ = 488 nm leads to d = 118 nm. If considering the largest angles (corresponding to the outer annulus radius,producing evanescent waves with the shortest decay length), the effective NA is NA_oBJ_ = 1.7, leading to d= 37 nm. By these simple calculations, the “average” decay thus lies between 37 nm – 118 nm, and is weighted by the distribution of intensity in the annular excitation.

Since such an intensity distribution is difficult to measure accurately, we instead opted to measure the average evanescent decay length more directly using silica beads (diameter 7.27 µm, refractive index, 1.42, Bangs Laboratories) placed in a solution of fluorescein dye (Fluka, Cat #32615) (**Supplementary Fig. 3a**). In this method, the known diameter of the bead is used to convert the apparent radii observed with TIRF to an axial depth^25^, z (**Supplementary Fig. 3b**). Following previous work^26^, we integrate the intensity I(z) from the coverslip surface to some depth z, as this corresponds to the observed signal F(z) at each depth. First, we assume the fluorescence is well modeled by a sum of two exponentials. The first term corresponds to signal derived from “pure” TIRF (with decay d) and the second term models scattering that is known to contaminate objective-type TIRF (with decay D): I(z) = Aexp(-z/d) + Bexp(-z/D),

where A and B are constants that account for incident beam intensity, concentration, and the relative weight of the scattering term. Integrating this expression yields

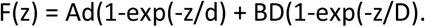

Fitting the measured fluorescence intensity at each depth (derived at each bead radius) to this expression (**Supplementary Fig. 3c**) with the MATLAB curve fitting toolbox gave d = 123 nm with 95% confidence interval (117nm, 129 nm). The scattering amplitude B represented ˜24 % of the signal.

### Diffraction limited TIRF imaging

Widefield TIRF images of GFP-HRas (**Supplementary Fig. 7a**) were acquired using a home-built system^27^ based on an olympus IX81 inverted microscope equipped with a 150X oil-immersion objective lens (olympus, N.A. = 1.45), a multi-band dichroic (405/488/561/633 BrightLine® quad-band bandpass filter, Semrock), and acousto-optic tunable filter (AoTF) and excitation lasers (405 nm, 488 nm, 561 nm and 633 nm, Coherent). GFP-HRas was excited at 488 nm laser with 100 ms exposures. Fluorescence was collected by an electron multiplying CCD (iXon888, Andor) after being filtered through amulti-bandemission0020filter(446/523/600/677 nm BrightLine® quad-band bandpass filter, Semrock). The microscope, AoTF and cameras were controlled through Micro-Manager^28^.

## Data Processing

### Deconvolution

Unless otherwise indicated, data presented in this paper were deconvolved to further enhance spatial resolution. Before deconvolution, background was subtracted from the raw images. Background was estimated by averaging 100 **"**dark″ images acquired without illumination. For deconvolution, we used the Richardson-Lucy algorithm^29,30^, blurring with a 2D PSF:

*For k* = 1, 2, … *N*

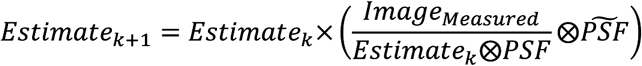

where ⊗ denotes convolution operation, *Image*_*Measured*_ is the measured image (after background subtraction) and 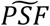is the flipped PSF:

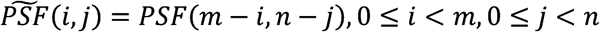

with *m, n* the PSF dimensions.

The PSF was experimentally derived by registering and then averaging the images of 20 100 nm yellow-green beads. Deconvolution was implemented in MATLAB 2017a with the number of iterations *N* set to 10.

### Tracking

For tracking the particles in the Rab11 dataset (**Fig. 3**), we performed semi-automated tracking using the TrackMate ImageJ Plugin^31^ (https://imagej.net/TrackMate). For particle detection, the Difference of Gaussian (DoG) detector was used with estimated blob diameter of 0.3 μm, and an initial quality threshold of 120. The particles were further filtered and linked with a simple Linear Assignment Problem (LAP) linker. Linking maximum distance and Gap-closing maximum distance were set to 0.3 μm, and the maximum frame gap was set to 5. For tracking on the whole image (**Fig. 3**), the linking filters were manually adjusted to filter out obviously spurious tracks. Then manual editing was performed within the plugin interface to improve tracking results. Within the cropped region used for downsampling analysis (**Supplemental Fig. 9, Supplemental Video 12**), images were downsampled 5- and 10 times in the time domain. Then, automated tracking was performed independently for the cropped images (100Hz) and the downsampled images (20 Hz and 10 Hz) without manually adjusting either linking filters or links. From the particle tracks (i.e., the sequences of coordinates denoting the position of each tracked particle at each time point), we computed several quantitative metrics including displacement, distance, instantaneous speed, mean speed and mean squared displacement (MSD). Given a trajectory consisting of *N* time points and the particle coordinates at *i*th time point ***p***_*i*_ = (*x*_*i*_, *y*_*i*_), we define the distance between any two points ***p***_*i*_ and ***p***_*j*_ as the Euclidean norm

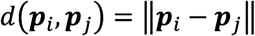

The total distance traversed at the *j*th time point is calculated from the starting point (the 1st time point) and defined as

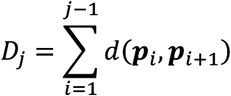

and the displacement (magnitude), also known as net distance

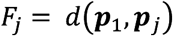

Then the total distance for the whole trajectory is *D*_*N*_ and the total displacement for thewholetrajectory is *F*_*N*_.

The instantaneous speed is defined as

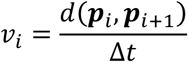

where Δ*t* is the time interval between two successive time points. The instantaneous speed is also the derivative of the traveled distance *D*_*j*_.

Then the mean speed is calculated as the average of the instantaneous speed:

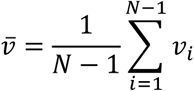

The mean squared displacement is calculated as

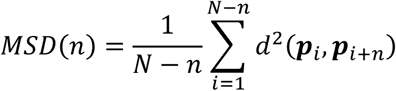

### Bleach correction

For several time-lapse datasets (**Fig. 2, 3, Supplementary Video 1, 2, 4, 10**), we performed standard bleaching correction using an ImageJ Plugin (Bleach Correction^32^, <u>https://imagej.net/Bleach_Correction</u>) with the “simpleratio” method.

### Flat fielding

Due to the spatially nonuniform profile of the excitation laser beam, the excitation intensity in both conventional- and TIRF-SIM is not distributed uniformly even when the excitation is scanned. The scanned excitation distribution has highest intensity in the center of the field of view and diminishes at increasing distances perpendicular to the scanning direction. In an attempt to normalize for this variation in excitation intensity (‘flat fielding’), in some of the datasets (**Fig. 2, 3**, **Supplementary Fig. 6, 8, 9, Supplementary Video 1, 2, 4-7, 11)** we averaged 100 images of a thin fluorescein layer, smoothed the average perpendicular to the scan direction, and divided the raw data by this smoothed average prior to deconvolution.

### Image display

All images are displayed in grayscale, except images in **Fig. 2 e-i**, displayed in green and/or magenta colormaps derived from ImageJ.

### Sample preparation

### Fixed Samples

For imaging microtubules within fixed samples (**Fig. 1, Supplementary Fig. 4**), high index coverslips were first immersed in 70% ethanol for ˜ 1 min and allowed to air dry in a sterile cell culture hood. U2oS cells were grown on uncoated high index coverslips until ˜50% confluency. The entire coverslip was submerged for 3 minutes in methanol pre-chilled to −20 °C to fix the cells. Coverslips were then washed in PBS at room temperature extensively before blocking in antibody dilution buffer (Abdil; 1%BSA, 0.3% Triton-X 100 in PBS) for 1 hour at room temperature. The primary antibody stain was performed overnight at 4 °C using 1/500 mg/ml of mouse anti α-Tubulin (Thermo Scientific #62204) in Abdil. The secondary antibody stain was performed for 1-2 hours at room temperature using 1/200 mg/ml of goat anti-mouse Alexa 488 (Invitrogen A11001) in Abdil.

### Live Jurkat T cells

Highindex coverslips were rinsed with 70% ethanol and dried with filtered air. The coverslips were then incubated in Poly-L-Lysine (PLL) at 0.01% W/V (Sigma Aldrich, St. Louis, Mo) for 10 min. PLL solution was aspirated and the coverslip was left to dry for 1 hour at 37 °C. Coverslips were next incubated with streptavidin (Invitrogen) at 2 μg/ml for 1 hour at 37°C and excess streptavidin was washed with PBS. Antibody coating for T cell activation was performed by incubating the coverslips in a 10 μg/ml solution of biotin labeled anti-CD3 antibody (oKt3, eBiosciences, San Diego, CA) for 2 hours at 37 °C. Excess antibody was removed by washing with L-15 imaging media immediately prior to the experiment. E6-1 Jurkat T-cells were transiently transfected with EMTB-3xEGFP (**Fig. 2a, b, Supplementary Video 1**) or F-tractin EGFP (**Supplementary Video 8**) plasmid using the Neon (Thermofisher Scientific) electroporation system two days before the experiment. Transfected cells were centrifuged and resuspended in L-15 imaging media prior to pipetting them onto the coverslip. Imaging was performed 10 minutes after the cells settled on the substrate. EMTB-3xEGFP was a gift from William Bement (Addgene plasmid # 26741) and pEGFP-C1 F-tractin EGFP was a gift from Dyche Mullins (Addgene plasmid # 58473).

### Live U2oS cells

### Ras, Rab, VSVG, ER imaging

Human osteosarcoma U2oS cells were routinely passaged in DMEM (Life technologies) plus 10% FBS (Hyclone) at 37 °C, with 5% Co2. For cleaning prior to live cell imaging, high index coverslips were boiled for 5 minutes with distilled water, thoroughly rinsed with distilled water and stored in 90% ethanol for at least 2 hours. In order to facilitate cell adherence, the coverslips were coated with FBS for 2 hours at 37°C. 24 - 48 hours prior to transfection, cells were plated on cleaned coverslips, at a density of ˜60%.Cells were transfected with the appropriate plasmid using Turbofect (Life Technologies) at a ratio of 3:1 (Liposomes:DNA). The next day, the medium was replaced with fresh DMEM plus 10% FBS without phenol red, which was also used as the imaging medium. To monitor wild type Ras dynamics, we used EGFP-HRas (**Fig. 2c, d, Supplementary Fig. 7, Supplementary Video 2, 3**), or if imaged with VSVG-GFP (Addgene #11912) (**Fig. 2e-g, Supplementary Video 4**), we used a HaloTag chimera of HRas (gift of Dominic Esposito, NCI). Halotag proteins were labeled using Janelia Fluor549 (gift of Luke Lavis, Janelia Research Campus) at a final concentration of 100 nM for 15 minutes. Following labelling, the cells were rinsed twice with plain DMEM, incubated with fresh medium plus 10% FBS for 20 minutes, and finally the medium replaced with fresh, phenol red free DMEM plus 10% FBS. The dynamics of Rab GTPase were followed for GFP tagged Rab11^33^ (Addgene #12674)(**Fig. 3, Supplementary Fig. 8, 9, Supplementary Video 11-13**). For dual labelling of Ras and the endoplasmic reticulum, we co-transfected the EGFP-HRas construct with pDsRed2-ER (Clontech, cat #632409) (**Fig. 2h, i, Supplementary Video 5**), which carries a KDEL ER retention signal.

### Myosin imaging

For imaging moysin IIA (**Supplementary Fig. 6a, Supplementary Video 10**), high index coverslips were plasma cleaned (PDC-001, Harrick Plasma) for 5 minutes, and then coated with 10 μg/ml human plasma fibronectin (Millipore, cat. # FC010) in PBS (ThermoFisher). U2oS cells were cultured in McCoys media (Invitrogen) supplemented with 10% fetal calf serum (ThermoFisher), at 37 °C in 5% Co_2_. Cells were transfected with GFP-myosin IIA expression and mApple-F-tractin plasmids as previously described^34^ and cultured for 12 hours prior to plating on fibronectin coated coverslips.

### Actin imaging

For actin imaging (**Supplementary Video 9**), U2oS cells were cultured at 37 °C in the presence of 5% Co2 in high glucose DMEM medium (ThermoFisher) with 10% fetal bovine serum,Pen/Strep and GlutaMAXTM (ThermoFisher). Cells were seeded on high index coverslips and transfected with Lifeact-GFP^35^ by X-tremeGENE HP (Sigma-Aldrich) 24 hours prior to imaging.

### Live INS-1 cells

For calcium imaging (**Supplementary Video 7**), INS-1 cells were cultured at 37°C with 5% Co_2_ in modified RPMI media (10% fetal bovine serum, 1% pen/strep, 11.1 mM glucose, 10 mM 4-(2-hydroxyethyl)-1-piperazineethanesulfonic acid [HEPES], 2 mM glutamine, 1 mM pyruvate, and 50 μM β-mercaptoethanol). Cells were seeded on high index coverslips that were cleaned by successive washing in detergent and bleach and then thoroughly rinsed with PBS. After PBS rinsing, coverslips with dipped in ethanol to sterilize and allowed to dry. Coverslips were treated with 0.1% poly-L-lysine (Sigma-Aldrich) for (5 to 10 minutes) followed by thorough rinsing with media. Cells were transfected with Lipofectamine 2000 (ThermoFisher) using 1 μg of DNA per coverslip. For calcium imaging, cells were transfected with GCamp6S-CAAX and imaged one day after transfection.

### Live SK-MEL cells

For FCHo (**Supplementary Fig. 6b**) imaging, SK-MEL cells were cultured in standard DMEM without phenol red (10% fetal bovine serum, 1% pen/strep, 1% GlutaMAX) at 37°C with 5% Co_2_. Cells were seeded and transfected (with FCHo2-GFP in this case) as described above for calcium imaging.

### Code Availability Statement

Deconvolution and simulation code are available from the corresponding author upon reasonable request.

### Data Availability Statement

The data that support the findings of this study are available from the corresponding author upon reasonable request.

## Supplementary Video Captions

**Supplementary Video 1.** Time-lapse imaging of Jurkat T cells expressing EMTB-3xEGFP at room temperature. Images were acquired every 50 ms, over 500 time points. Images were binned 2×2 relative to the data for display purposes. See also **Fig. 2a-b.**

**upplementary Video 2.** Time-lapse imaging of U2oS cell expressing EGFP-HRas at 37 °C. Images were acquired every 0.75 s, over 60 time points. Images were binned 2×2 relative to the data for display purposes. See also **Fig. 2c.**

**Supplementary Video 3.** Higher magnification view of subregion in **Supplementary Video 2,** highlighting the ‘wave-like’ dynamics among Ras microdomains. See also **Fig. 2d.**

**Supplementary Video 4.** Dual-color time-lapse imaging of U2oS cell expressing EGFP-VSVG (green) and Halotag-Ras (magenta) at 37 °C. Images were acquired every 2.3 s, over 100 time points. Images were binned 2×2 relative to the data for display purposes. See also **Fig. 2e-g.**

**Supplementary Video 5.** Dual-color (merge, bottom) time-lapse imaging of U2oS cell expressing EGFP-HRas (green, middle) and pDsRed2-ER (magenta, top) at 37 °C. Images were acquired every 1.2 s, over 100 time points. Images were binned 2×2 relative to the data for display purposes. See also **Fig. 2h.**

**Supplementary Video 6.** Higher magnification view of subregion in **Supplementary Video 5,** highlighting apparent fission and fusion of Ras microclusters. See also **Fig. 2i.**

**Supplementary Video 7.** Time-lapse imaging of calcium flux within INS-1 cells, as reported by GCamp6S-CAAX at room temperature. Images were acquired every 88 ms, over 500 time points. Note localized calcium activity. Images were binned 2×2 relative to the data for display purposes.

**Supplementary Video 8.** Time-lapse imaging of Jurkat T cells expressing F-tractin EGFP at room temperature. Images were acquired every 2 s, over 200 time points.

**Supplementary Video 9.** Time-lapse imaging of U2oS cell expressing Lifeact-GFP at 37 °C. Images were acquired every 60 ms, over 50 time points.

**Suppleentary Video 10.** Dual-color time-lapse imaging of U2oS cell expressing GFP-myosin IIA (green) and mApple-F-tractin (magenta) at 37 °C. Images were acquired every 6 s, over 50 time points.

**Supplementary Video 11.** Time-lapse imaging of U2oS cell expressing GFP-Rab11 at 37 °C. Images were acquired at 100Hz frame rate, over 600 time points. Images were binned 2×2 relative to the data for display purposes. See also **Fig. 3.**

**Supplementary Video 12.** Higher magnification view of subregion in **Supplementary Video 11** highlighting diffusive (blue) vs. directed (red) motion of particles. The first 305 frames (from 0 s to 3.04s) are shown. See also **Fig. 3 b-d.**

**Supplementary Video 13.** Higher magnification view of subregion in **Supplementary Video 11** highlighting tracks derived from many particles. The 300^th^ to 500^th^ frames from the total image series (from 3 s to 5 s) are shown. See also **Supplementary Fig. 9.**

## References

1 Poulter, N. S., Pitkeathly, W. T. E., Smith, P. J. & Rappoport, J. Z. in *Advanced Fluorescence Microscopy: Methods and Protocols* Vol. 1251 Methods in Molecular Biology (ed Peter J Verveer) (Springer, 2015).

2 Fiolka, R., Beck, M. & Stemmer, A. Structured illumination in total internal reflection fluorescence microscopy using a spatial light modulator. optics Letters 33, 1629–1631 (2008).

3 Kner, P., Chhun, B. B., Griffis, E. R., Winoto, L. & Gustafsson, M. G. L. Super-resolution video microscopy of live cells by structured illumination. Nat. Methods 6, 339–342 (2009).

4 Chung, E., Kim, D., Cui, Y., Kim, Y. H. & So, P. T. Two-dimensional standing wave total internal reflection fluorescence microscopy: superresolution imaging of single molecular and biological specimens. Biophys. J. 93, 1747–1757 (2007).

5 Gliko, o., Reddy, G. D., Anvari, B., Brownell, W. E. & Saggau, P. Standing wave total internal reflection fluorescence microscopy to measure the size of nanostructures in living cells. J Biomed opt 11, 064013 (2006).

6 Li, D. et al. Extended-resolution structured illumination imaging of endocytic and cytoskeletal dynamics. Science 349, aab 3500 (2015).

7 York, A. G. et al. Instant super-resolution imaging in live cells and embryos via analog image processing. Nat Methods 10, 1122–1126 (2013).

8 Roth, S., Sheppard, C. J. R., Wicker, K. & Heintzmann, R. optical photon reassigment microscopy (oPRA). optical Nanoscopy 2, 1–6 (2013).

9 De Luca, G. M. et al. Re-scan confocal microscopy: scanning twice for better resolution. Biomed opt Express 4, 2644–2656 (2013).

10 Curd, A. et al. Construction of an instant structured illumination microscope. Methods 15, 30029–30023 (2015).

11 Stout, A. L. & Axelrod, D. Evanescent field excitation of fluorescence by epi-illumination microscopy. Applied optics 28, 5237–5242 (1989).

12 Gould, T. J., Myers, J. R. & Bewersdorf, J. Total internal reflection STED microscopy. opt Express19, 13351–13357 (2011).

13 Burnette, D. T. et al. A contractile and counterbalancing adhesion system controls the 3D shape of crawling cells. J Cell Biol. 205, 83–96 (2014).

14 Sochacki, K. A., Dickey, A. M., Strub, M. P. & Taraska, J. W. Endocytic proteins are partitioned at the edge of the clathrin lattice in mammalian cells. Nat Cell Biol 19, 352–361 (2017).

15 Faire, K. et al. E-MAP-115 (ensconsin) associates dynamically with microtubules in vivo and is not a physiological modulator of microtubule dynamics. J Cell Sci 112, 4243–4255 (1999).

16 Miller, A. L. & Bement, W. M. Regulation of cytokinesis by Rho GTPase flux *Nat Cell Biol* 11, 71–77 (2009).

17 Apolloni, A., Prior, I. A., Lindsay, M., Parton, R. G. & Hancock, J. F. H-ras but Not K-ras Traffics to the Plasma Membrane through the Exocytic Pathway. Mol Cell Biol. 20, 2475–2487 (2000).

18 Grimm, J. B. et al. A general method to improve fluorophores for live-cell and single-molecule microscopy. Nat Methods 12, 244–250 (2015).

19 Presley, J. F. et al. ER-to-Golgi transport visualized in living cells. Nature 389, 81–85 (1997).

20 Choy, E. et al. Endomembrane Trafficking of Ras: The CAAX Motif Targets Proteins to the ER and Golgi. Cell 98, 69–80 (1999).

21 Takahashi, S. et al. Rab11 regulates exocytosis of recycling vesicles at the plasma membrane. J Cell Sci 125, 4049–4057 (2012).

22 Fiolka, R. Clearer view for TIRF and oblique illumination microscopy. optics Express 24, 29556–29567 (2016).

23 olveczky, B. P., Periasamy, N. & Verkman, A. S. Mapping Fluorophore Distributions in Three Dimensions by Quantitative Multiple Angle-Total Internal Reflection Fluorescence Microscopy. Biophys. J. 73, 2836–2847 (1997).

24 Winter, P. W. et al. Two-photon instant structured illumination microscopy improves the depth penetration of super-resolution imaging in thick scattering samples. optica 1, 181–191 (2014).

25 Mattheyses, A. L. & Axelrod, D. Direct measurement of the evanescent field profile produced by objective-based total internal reflection fluorescence. Journal of Biomedical optics 11, 014006 (2006).

26 Fu, Y. et al. Axial superresolution via multiangle TIRF microscopy with sequential imaging and photobleaching. Proc Natl Acad Sci U S A 113, 4368–4373 (2016).

27 York, A. G., Ghitani, A., Vaziri, A., Davidson, M. W. & Shroff, H. Confined activation and subdiffractive localization enables whole-cell PALM with genetically expressed probes. Nature Methods 8, 327–333 (2011).

28 Edelstein, A. D. et al. Advanced methods of microscope control using **μ**Manager software. Journal of Biological Methods 1, e11 (2014).

29 Richardson, W. H. Bayesian-Based Iterative Method of Image Restoration. JoSA 62, 55–59 (1972).

30 Lucy, L. B. An iterative technique for the rectification of observed distributions. Astronomical Journal 79, 745–754 (1974).

31 Tinevez, J. Y. et al. TrackMate: An open and extensible platform for single-particle tracking. Methods 115, 80–90 (2017).

32 Miura, K., Rueden, C., Hiner, M., Schindelin, J. & Rietdorf, J. ImageJ Plugin CorrectBleach V2.0.2. Zenodo (2014).

33 Choudhury, A. et al. Rab proteins mediate Golgi transport of caveola-internalized glycosphingolipids and correct lipid trafficking in Niemann-Pick C cells. J Clin Invest 109, 1541–1550 (2002).

34 Baird, M. A. et al. Local pulsatile contractions are an intrinsic property of the myosin 2A motor in the cortical cytoskeleton of adherent cells. Mol Biol Cell 28, 240–251 (2017).

35 Riedl, J. et al. Lifeact: a versatile marker to visulize F-actin. Nat Methods 5, 605–607 (2008)

